# A minimally-invasive method for ancient DNA sampling of Prehistoric bone and antler tools and hunting weapons

**DOI:** 10.1101/2023.04.02.535282

**Authors:** José-Miguel Tejero, Olivia Cheronet, Pere Gelabert, Brina Zagorc, Esteban Alvarez, Aline Averbouh, Guy Bar-Oz, Anna Belfer-Cohen, Marjolein D. Bosch, Florian Brück, Marián Cueto, Martin Dockner, Josep Maria Fullola, Diego Gárate, Michael Giannakoulis, Cynthia González, Nino Jakeli, Xavier Mangado, Tengiz Meshveliani, Petr Neruda, Philip Nigst, Petra G. Šimková, Jesús Tapia, Marta Sánchez de la Torre, Catherine Schwab, Gerhard Weber, Ron Pinhasi

## Abstract

Internal and external bony tissues from diverse mammalian taxa are one of the primary animal raw materials exploited for technical and symbolic purposes by Eurasian Upper Palaeolithic hunter-gatherers. Identifying the source species used for osseous raw material is critical to gain insights into these populations’ behaviour, technology, and subsistence. The study of osseous tools has advanced in the last few years by combining archaeological and biomolecular methods. Ancient genomics opens many new analytical opportunities. Ancient DNA (aDNA) can provide a wealth of information about the animal sources of these objects. Unfortunately, aDNA analyses often involve destructive sampling. Here, we develop and apply a minimally-invasive aDNA sampling method for an assemblage of 42 prehistoric hunting weapons and tools from various Eurasian archaeological sites. We evaluated the impact of our approach on the specimens visually, microscopically and through Micro-CT scans. The surface impacts are marginal, ranging from 0.3-0.4 mm. Using a custom-made DNA capture kit for 54 mammalian species, we obtained sufficient aDNA to identify the taxa of 33% of the objects. For one of the tools, we recovered enough endogenous aDNA to infer the genetic affinities of the individual. Our results also demonstrate that ancient antler, one of the primary raw materials used during a large part of prehistory, is a reliable source of aDNA. Our minimally-invasive aDNA sampling method is therefore effective while preserving osseous objects for potential further analyses: morphometric, technical, genetic, radiometric and more.

## Introduction

Objects made from diverse internal and external skeletal tissues (e.g., bone, antler, ivory, teeth) are one of the most common archaeological remains recovered at prehistoric sites from the Palaeolithic to recent periods. The worked osseous raw materials were sourced from a wide range of vertebrate species. The variable morpho-structural properties of the raw materials constrain the technical possibilities to exploit them. Identifying the species of the skeletal raw material is critical to gain insights into how this selection fits into a given environment and cultural system and to understand how these populations exploited their environment (economic aspects), how they saw themselves within this environment (social and symbolic aspects) and how they transformed it (technological aspects) (Bradfield et al., 2021; Langley et al., 2020; Martisius, McPherron, et al., 2020; Pedergnana et al., 2021; Sidéra, 2000; Tejero et al., 2018, 2021).

For the majority of osseous objects, the designation of raw material by macroscopic analysis is restricted to bone, antler and ivory (Pacher, 2010). Categorising osseous tissues’ taxonomic origin is generally only possible using biomolecular methods, albeit some attempts by X-Ray micro-tomography have been made (Lefebvre et al., 2016) to differentiate between red deer and reindeer antlers. A major difficulty lies in the identification of the intensely transformed anatomical blank during the objects’ production, involving the loss of many, if not all, specific diagnostic attributes. Nevertheless, despite the importance of the diverse skeletal tissues for (pre)historic past societies, palaeogenetics and palaeoproteomics of osseous objects analyses have mainly focused on bone artefacts (Bradfield et al., 2021; Martisius, Welker, et al., 2020; McGrath et al., 2019; Pacher & Hofreiter, 2004). Genetic studies of other raw materials, such as antler, are mostly restricted to modern specimens in the context of deer conservation (e.g., (Bi et al., 2020; Greco et al., 2021; Hoffmann et al., 2015; Venegas et al., 2020)), with a single palaeontological Giant dear (*Megaloceros giganteus*) example from an unclear context with an estimated age of around 12,000 years (Kuehn et al., 2005). Ancient DNA (aDNA) analyses, sometimes in combination with palaeoproteomics, of deer antlers from archaeological contexts have been restricted to recent prehistoric periods (pre-Viking contexts from Scotland and Scandinavia (von Holstein et al., 2014) and Medieval times (Rosvold et al., 2019)). Unlike bone, where this has now been extensively tested, the long-term ability of antler to preserve DNA therefore remains unclear.

Here we present the results of the first aDNA analysis of a diverse set of 42 antler and bone hunting weapons, including also some domestic tools from Palaeolithic and Neolithic sites (c. 39-8 ka) from South-Western Europe (France, Spain), Central Europe (Austria, Czech Republic), the Caucasus (Georgia), and the Levant (Lebanon, Israel). We present and utilise a minimally-invasive aDNA sampling method originally developed for human teeth (Harney et al., 2021) that we optimised for osseous objects. The method is combined with a new custom-created set of capture baits for the mitochondrial DNA of 54 mammalian species. The obtained mitochondrial data can be used for identifying the exploited taxa and potentially the phylogeny of the individual. Our study also establishes that ancient antler is a reliable source of aDNA, which is particularly relevant considering its prevalent use throughout the Upper Palaeolithic, Mesolithic and later periods. We quantitatively assess the invasiveness of our new method on the objects by studying both their macro-morphology, and their structure. Macroscopic and microscopic assessments as well as micro-CT scans confirmed that the macro-morphology of objects remains broadly unchanged after sampling. This allows for a range of further studies to be carried out on the objects after sampling, including morphometric, technical, genetic, radiometric and more analyses.

## Material and methods

The analysed assemblage comprises 42 Upper Palaeolithic items encompassing hunting weapons (projectile points and one harpoon), blanks, production wastes and domestic tools (awls) (Table 1). We recorded and sampled one specimen from the Aurignacian layers (S-III, A⍵) of Isturitz (France) (Normand et al., 2007); one from the Early Aurignacian layer from La Quina-Aval (France) (Dujardin, 2001; G. Henri-Martin, 1958, 1965; L. Henri-Martin, 1930); one from the Aurignacian layers of La Ferrassie (France); two from the Early Aurignacian layer (I) of Abri Poisson (France) (Peyrony, 1932); five specimens from Aurignacian layers of Mladeč cave (Czech Republic) (Teschler-Nicola, 2006); two examples from the Upper Palaeolithic and Epipalaeolithic layers of Ksâr ‘Akil (Lebanon) (Bosch et al., 2015; Ewing, 1948; Newcomer, 1974); three items from the Magdalenian layers (UE103) of Tito Bustillo (Spain) (Álvarez-Fernández, Esteban Tapia, Jesús. Agirre-Uribesalgo, Amaia Arias, Pablo Camarós, Edgard Cerezo-Fernández, Rosana García Alonso, Beatriz Martín-García, Noelia Martín-Jarque, Sergio Peyroteo-Stjerna, Rita Portero, Rodrigo Teira, L.uis C. Cueto, Marián, 2022); four objects from the Magdalenian layers (201, 301, 305) of Cueva Chufín (Spain) (Cabrera Valdés, 1977), nine samples from the Upper Palaeolithic (Unit C) of Dzudzuana (Georgia) (Bar-Yosef et al., 2011); seven items from Satsurblia (Georgia) Upper Palaeolithic layers (BIII, BIV) with three additional experimental ones made from unmodified faunal remains (fragments of long mammal bones from layer AIIb) (Pinhasi et al., 2014; Tejero et al., 2021); one Neolithic item from Samele Klde (Georgia); 2 items from Nahal Rahaf (Israel) Arqov-Dishon layers (Layers 5, and 7b) (Barzilai et al., 2020; Shemer et al., 2023); and four pieces from the Late and Middle Magdalenian layers (NII-NIII) of Cova del Parco (Spain) (Tejero, 2005; Tejero & Fullola, 2008).

**Table 1:**
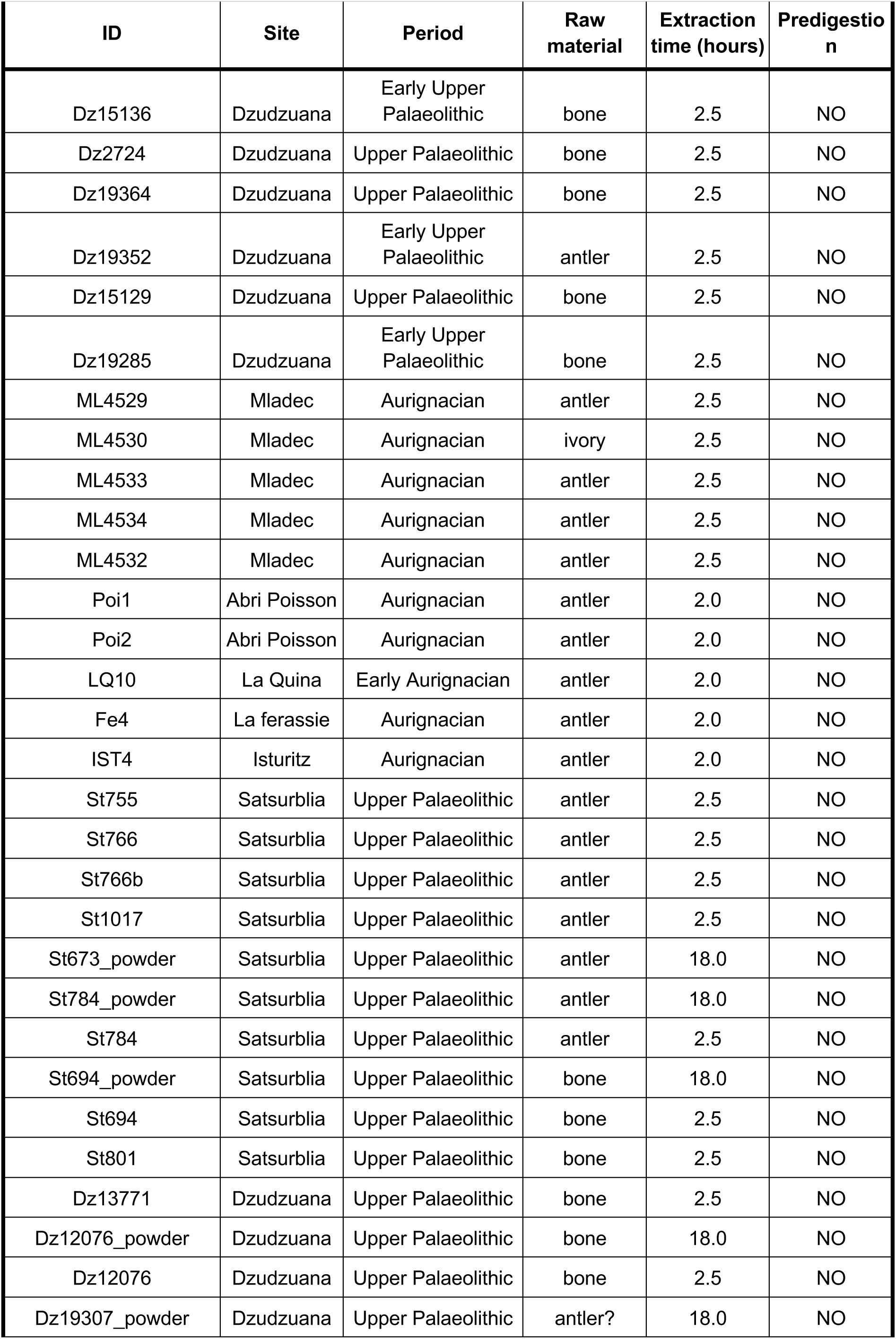
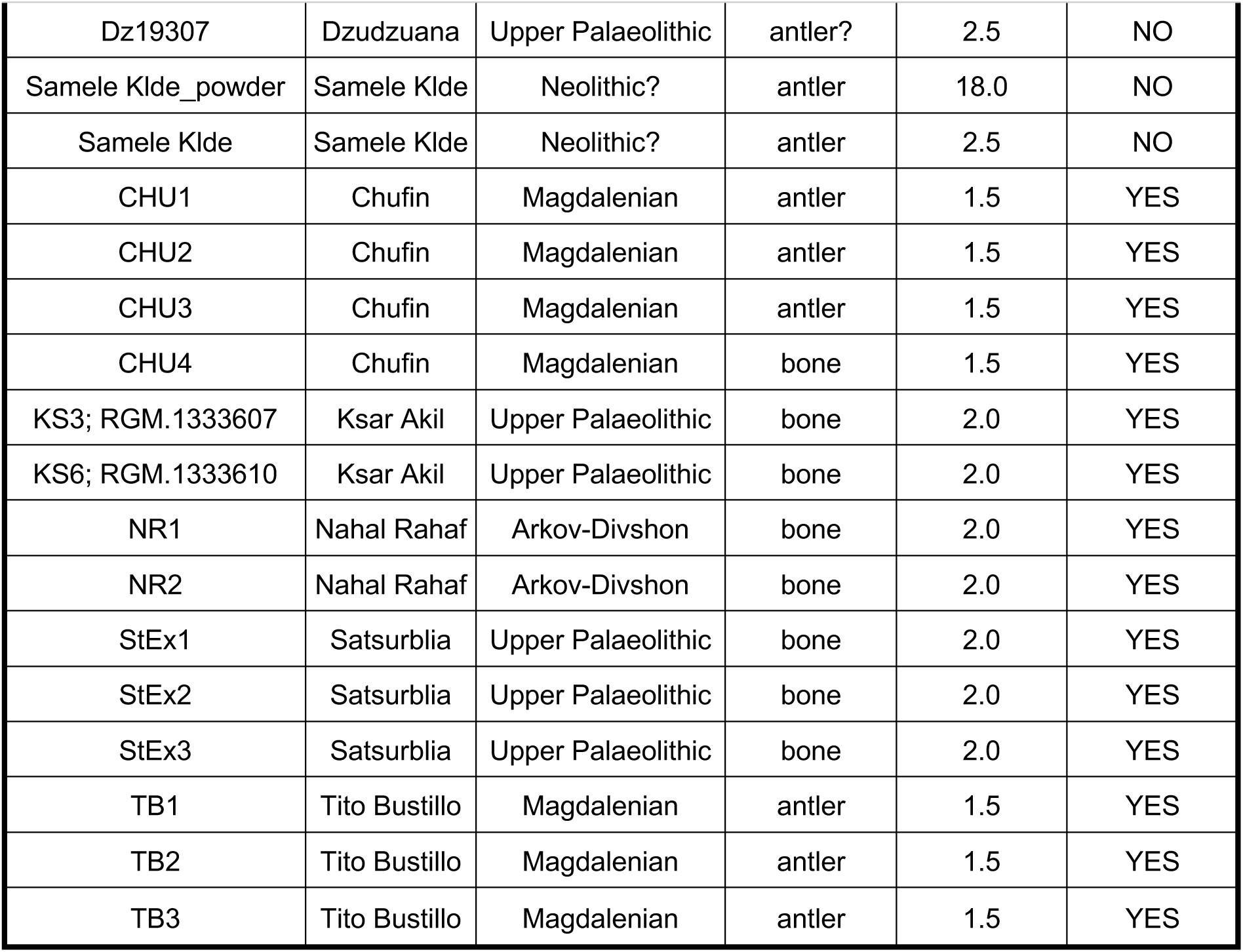
Description of the samples and sampling conditions.

| ID | Site | Period | Raw material | Extraction time (hours) | Predigestion |
| --- | --- | --- | --- | --- | --- |
| Dz15136 | Dzudzuana | Early Upper Palaeolithic | bone | 2.5 | NO |
| Dz2724 | Dzudzuana | Upper Palaeolithic | bone | 2.5 | NO |
| Dz19364 | Dzudzuana | Upper Palaeolithic | bone | 2.5 | NO |
| Dz19352 | Dzudzuana | Early Upper Palaeolithic | antler | 2.5 | NO |
| Dz15129 | Dzudzuana | Upper Palaeolithic | bone | 2.5 | NO |
| Dz19285 | Dzudzuana | Early Upper Palaeolithic | bone | 2.5 | NO |
| ML4529 | Mladec | Aurignacian | antler | 2.5 | NO |
| ML4530 | Mladec | Aurignacian | ivory | 2.5 | NO |
| ML4533 | Mladec | Aurignacian | antler | 2.5 | NO |
| ML4534 | Mladec | Aurignacian | antler | 2.5 | NO |
| ML4532 | Mladec | Aurignacian | antler | 2.5 | NO |
| Poi1 | Abri Poisson | Aurignacian | antler | 2.0 | NO |
| Poi2 | Abri Poisson | Aurignacian | antler | 2.0 | NO |
| LQ10 | La Quina | Early Aurignacian | antler | 2.0 | NO |
| Fe4 | La ferassie | Aurignacian | antler | 2.0 | NO |
| IST4 | Isturitz | Aurignacian | antler | 2.0 | NO |
| St755 | Satsurblia | Upper Palaeolithic | antler | 2.5 | NO |
| St766 | Satsurblia | Upper Palaeolithic | antler | 2.5 | NO |
| St766b | Satsurblia | Upper Palaeolithic | antler | 2.5 | NO |
| St1017 | Satsurblia | Upper Palaeolithic | antler | 2.5 | NO |
| St673_powder | Satsurblia | Upper Palaeolithic | antler | 18.0 | NO |
| St784_powder | Satsurblia | Upper Palaeolithic | antler | 18.0 | NO |
| St784 | Satsurblia | Upper Palaeolithic | antler | 2.5 | NO |
| St694_powder | Satsurblia | Upper Palaeolithic | bone | 18.0 | NO |
| St694 | Satsurblia | Upper Palaeolithic | bone | 2.5 | NO |
| St801 | Satsurblia | Upper Palaeolithic | bone | 2.5 | NO |
| Dz13771 | Dzudzuana | Upper Palaeolithic | bone | 2.5 | NO |
| Dz12076_powder | Dzudzuana | Upper Palaeolithic | bone | 18.0 | NO |
| Dz12076 | Dzudzuana | Upper Palaeolithic | bone | 2.5 | NO |
| Dz19307_powder | Dzudzuana | Upper Palaeolithic | antler? | 18.0 | NO |
| Dz19307 | Dzudzuana | Upper Palaeolithic | antler? | 2.5 | NO |
| Samele Klde_powder | Samele Klde | Neolithic? | antler | 18.0 | NO |
| Samele Klde | Samele Klde | Neolithic? | antler | 2.5 | NO |
| CHU1 | Chufin | Magdalenian | antler | 1.5 | YES |
| CHU2 | Chufin | Magdalenian | antler | 1.5 | YES |
| CHU3 | Chufin | Magdalenian | antler | 1.5 | YES |
| CHU4 | Chufin | Magdalenian | bone | 1.5 | YES |
| KS3; RGM.1333607 | Ksar Akil | Upper Palaeolithic | bone | 2.0 | YES |
| KS6; RGM.1333610 | Ksar Akil | Upper Palaeolithic | bone | 2.0 | YES |
| NR1 | Nahal Rahaf | Arkof-Divshon | bone | 2.0 | YES |
| NR2 | Nahal Rahaf | Arkof-Divshon | bone | 2.0 | YES |
| StEx1 | Satsurblia | Upper Palaeolithic | bone | 2.0 | YES |
| StEx2 | Satsurblia | Upper Palaeolithic | bone | 2.0 | YES |
| StEx3 | Satsurblia | Upper Palaeolithic | bone | 2.0 | YES |
| TB1 | Tito Bustillo | Magdalenian | antler | 1.5 | YES |
| TB2 | Tito Bustillo | Magdalenian | antler | 1.5 | YES |
| TB3 | Tito Bustillo | Magdalenian | antler | 1.5 | YES |

### Ancient DNA

DNA sampling was performed using two methods. For some pieces, a drill was used to collect ∼50 mg of powder from the object’s interior. The DNA was then extracted from powder following the protocol outlined by (Dabney et al., 2013) with modifications described in (Korlević et al., 2015), namely the replacement of the Qiagen Minelute column custom constructions for DNA purification with columns from the Roche High Pure Viral Nucleic Acid kitt. Most items were sampled using the new minimally-destructive extraction procedure presented here. It is based on the protocol described by (Harney et al., 2021) with a number of modifications detailed below.

The extractions were performed at the location of sample storage. The environment in which it was performed was cleaned as thoroughly as possible: surfaces were wiped with a dilute (about 1.2%) bleach solution and covered with a bleach-cleaned aluminium foil. In all cases, we first verified that no PCR was ever performed in the same space to avoid potential contamination.

The first step consisted in cleaning each object by first wiping with a bleach solution (about 1.2%) and subsequently rinsing thoroughly with absolute ethanol. The pieces were then exposed to short-wave UV light for 10 minutes on each surface.

Unlike the procedure described in the (Harney et al., 2021) protocol, the samples were not wrapped in Parafilm, but instead entirely submerged in extraction buffer. The exception was the samples stored at the Musée d’Archéologie Nationale (France), where the pieces were entirely wrapped in parafilm except for leaving a small window exposed (∼2-4 cm^2^). The smallest possible container was selected to fit the whole piece comfortably with as little spare space as possible. The possible containers were the following: 5 ml, 15 ml and 50 ml Eppendorf DNA LoBind tubes and sterile plastic bags.

In some cases, the object was wholly submerged for 20 minutes in extraction buffer, for a pre-digestion. The initial lysate was then discarded to remove the potential external DNA contamination. This was only performed for the later batch of samples containing the items from Chufín, Tito Bustillo Nahal Rahaf and Satsurblia (experimental items). The items were then re-submerged in extraction buffer, the volume of which was adapted for each piece. The minimum amount that enabled the pieces to be fully submerged ranged between 1.0 and 15.0 ml. The extraction was performed in room-temperature to warm conditions at ∼35 degrees C, with the liquid in the tubes moved around gently at regular 15-minute intervals, while monitoring the effect of the buffer on the piece’s surface condition. The duration of the extraction was adapted for each item. In all cases it was stopped as soon as any effect of digestion on the piece became visible. Individual digestion times are given in table 1, ranging from 0.5 to 3 hours.

The obtained lysates were cleaned up following (Dabney et al., 2013) with the same above-described modifications. As most samples resulted in more than 1 ml of extraction buffer, a ratio of 13:1 was used to calculate the amount of binding buffer required for optimal binding of the DNA to the silica columns, and the entire mixture run through the same column.

Subsequently, double-stranded libraries were built from 25.0 ul of extract, according to Meyer and Kircher (2010). Qiagen MinElute PCR Purification kits were used for the intermediate clean-up steps. The libraries were double-indexed and amplified using the NebNext Q5U Master Mix DNA Polymerase (NEB) using a number of cycles calculated using the qPCR analysis of 1 ul of library. Indexed libraries were captured using a custom build capture kit for the mitochondrial DNA of 54 mammalian species. This capture kit has been designed by the team in Vienna and produced by myBaits (Arbor Biosciences) (Supp. Table 1). The usage of this capture kit allows to screen for an extended list of species at the same time, both extending the possibilities to recover aDNA and also improving the discrimination capabilities, allowing species-specific hits and better discriminating between species from the same family. This was then shallow-sequenced as part of a larger pool of samples on a single lane of a NovaSeq SP system using the XP workflow.

### Bioinformatics

Sequenced reads were processed after demultiplexing. Sequenced adapters and short reads below 30 were discarded using Cutadapt 4.2 (Martin, 2011). The remaining reads were aligned against 40 representative mammalian species in a competitive mapping (list) with bwa aln 0.7.17 (Li & Durbin, 2009), disabling seeding and with a gap penalty open of 2. The aligned reads were filtered by quality with samtools 1.16.1 (Li et al., 2009), setting minimum mapping quality of 30 and removing duplicates with picard-tools 2.27.5 (*Picard-Tools*, n.d.). The remaining reads were inspected with mapdamage 2-2.2.1 (Jónsson et al., 2013) to determine the deamination patterns and with qualimap 2.2.1 (Okonechnikov et al., 2016) to inspect the results of the competitive mapping. Non-human species were considered positively identified when more than 50 reads could be mapped to the genome of a particular species. When more than one hit was present per sample, we focused on the dominant taxon (the one with the most mapped reads). We therefore considered this as the source.

Only samples which yielded more than 500 mammalian aDNA reads were further analysed. For these, we generated a consensus sequence with bcftools and vcfutils (Li et al., 2009). The consensus sequences were aligned with other present-day and modern animal sequences with Clustal Omega 1.2.4 (Sievers & Higgins, 2014), and we then performed a Maximum likelihood tree with the alignment using MEGA 10.2.4 (Tamura et al., 2013) with partial deletion and 100 bootstrap replications. All trees were plotted with MEGA.

### Macro and micro morphological analysis of objects

A comprehensive technical analysis of some of the pieces studied (Isturitz, La Quina, Abri Poisson, Satsurblia, Dzudzuana and Ksâr ‘Akil) was performed before and after the DNA extraction independently. Technical analysis is based on the assessment of the operational sequence and follows several distinct steps (Averbouh, 2000, 2001), including the identification of manufacture and use wear marks.

In order to identify the magnitude of damage caused by the extraction process, the items were scanned at the Vienna µCT Lab before and after extraction using an industrial Viscom X8060 NDT scanner (scanning parameters: 110-140 kV, 280-410 mA, 1400-2000 ms, 0.75 mm copper filter with a voxel size 23 µm). To obtain 3D surface models of the item, virtual segmentation of the µCT data was performed with Amira software (www.thermofisher.com). To further determine possible changes caused by our extraction method, we analysed the surfaces in Geomagic Design X 64 (www.3dsystems.com). 3D models were aligned according to homologous landmarks set on sufficiently visible morphological structures of both surfaces, and differences were assessed.

## Results

We captured mitochondrial DNA from 42 bone tools. For 20 of those (48%), we were not able to identify any non-human mammalian mitochondrial DNA. For seven samples (16%), the species identified contradicted the preliminary visual analysis, as it was determined that the items were made of antler, but the genetically identified species did not possess antlers (*Sus scrofa*, *Bos taurus*, and *Capra hircus*). These results can be explained by the conservation of the items in the museum. These three species are consistent with those used to make animal-based glues, commonly used in museum conservation (Schellmann, 2007). Finally, for 14 of the pieces (33%), it was possible to confidently identify the source species. While most of these yielded a very low mitochondrial coverage (<12x), a few yielded more data, one of which (CHU4) yielded a 176.91x *Cervus elaphus* mitochondrial genome, enabling further phylogenetic analyses. All results are summarised in table 2.

**Table 2:**
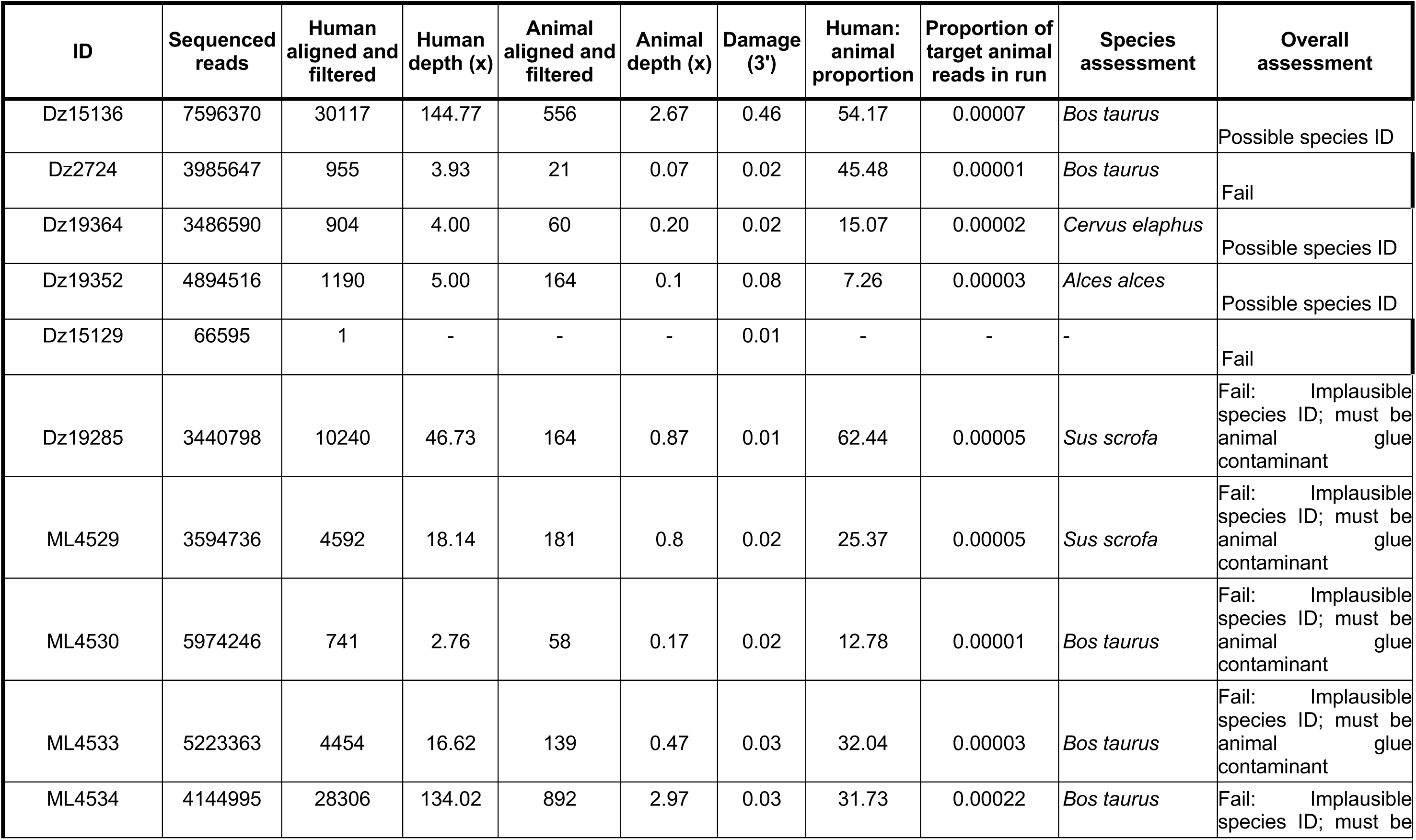
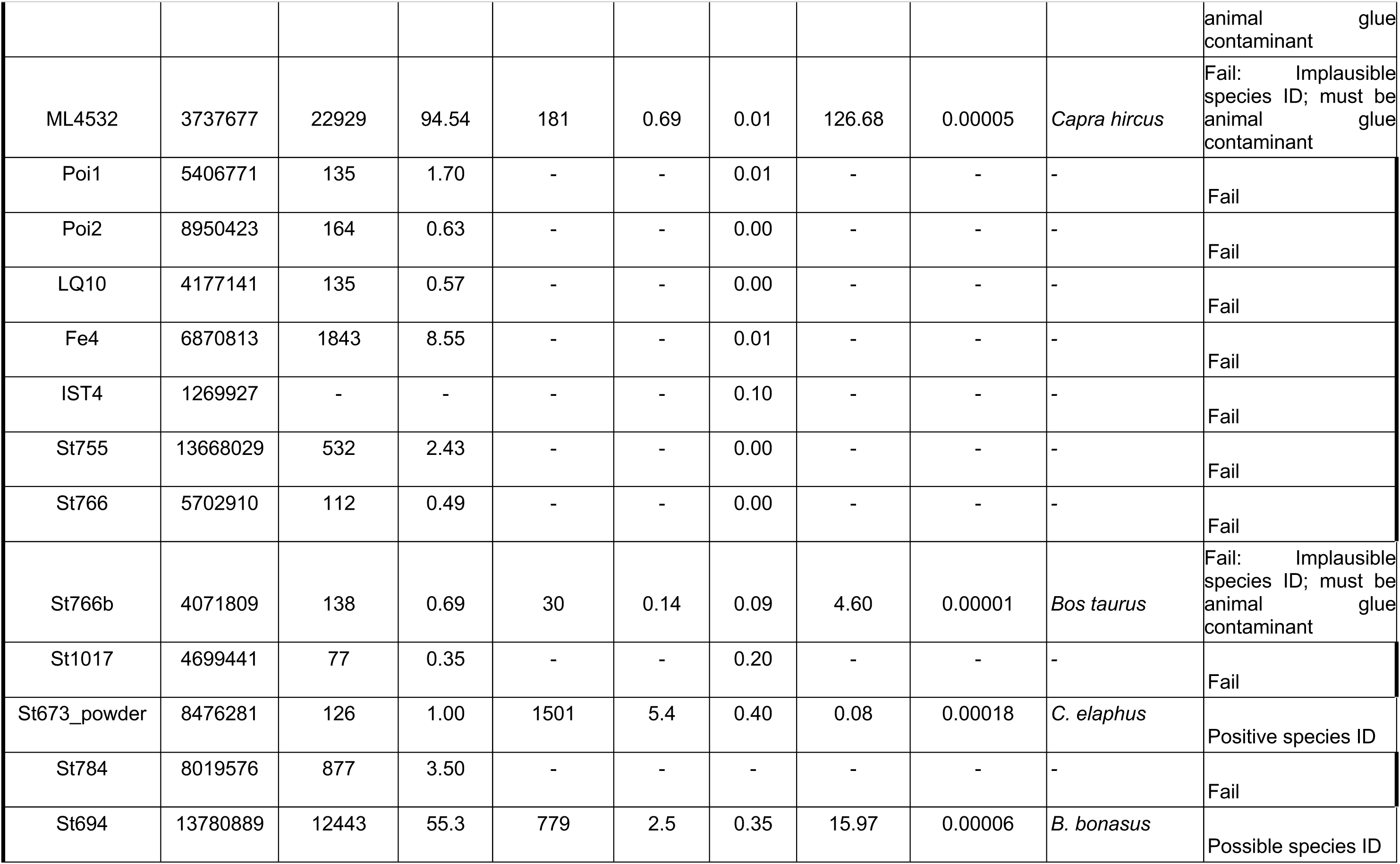
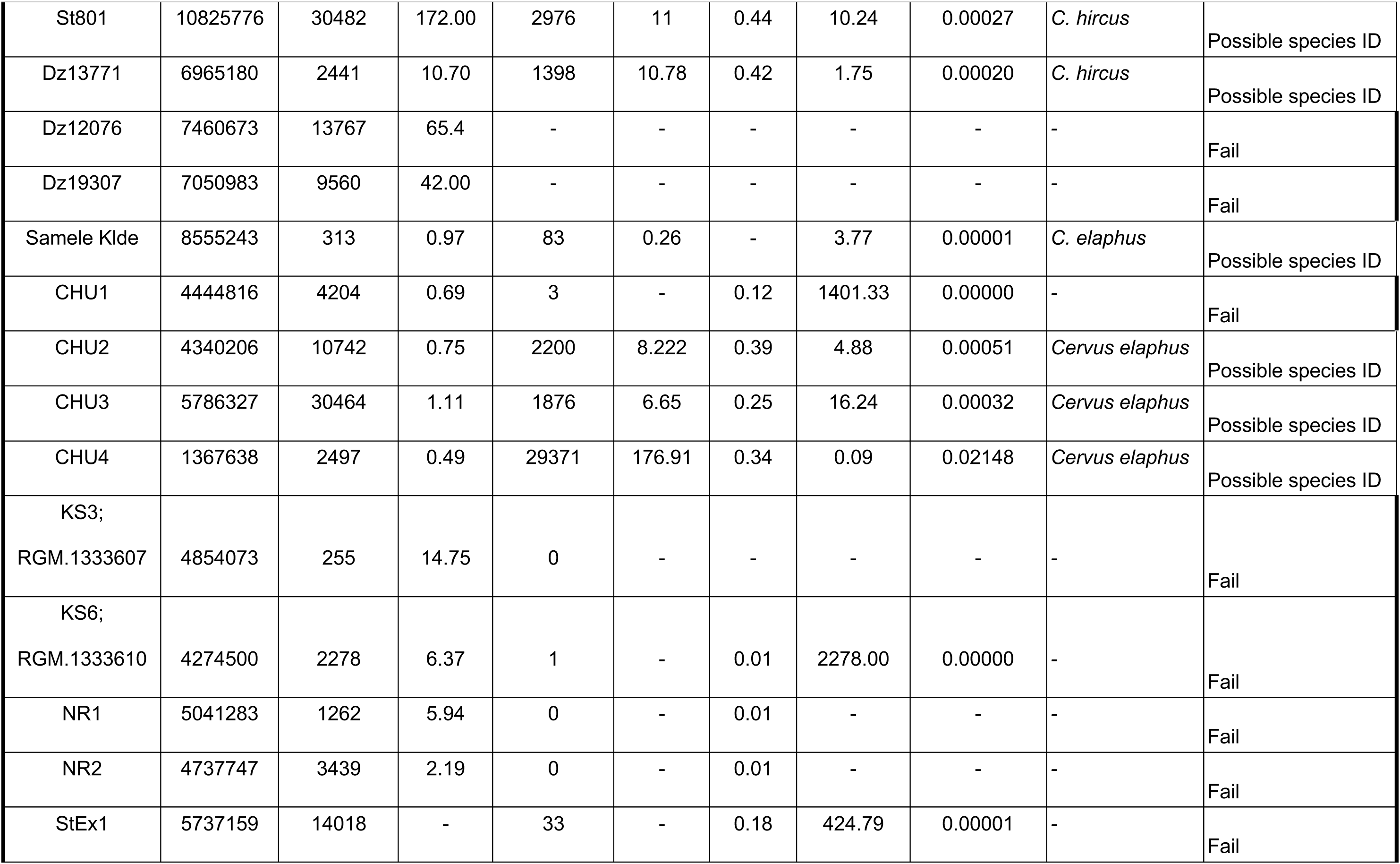
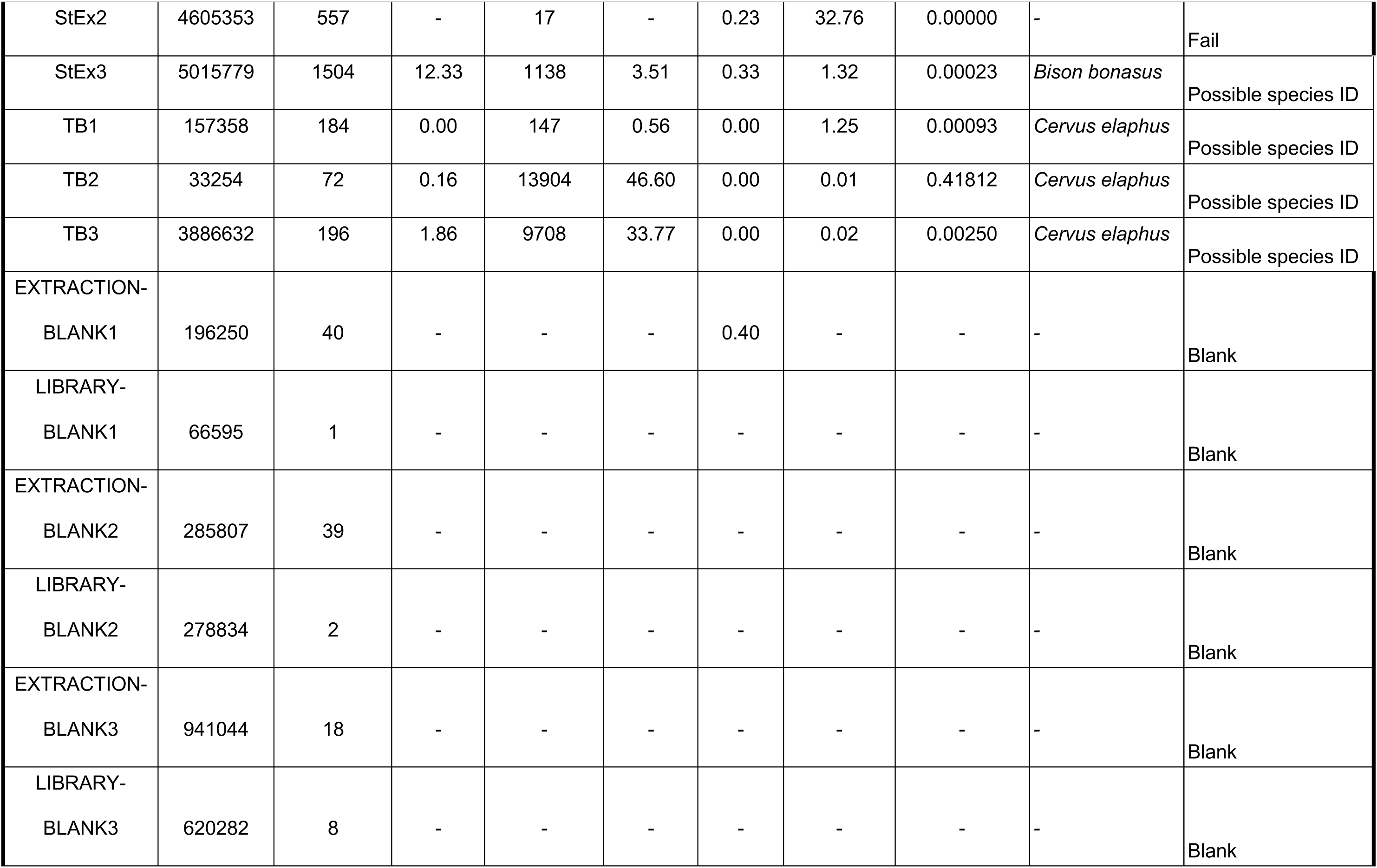
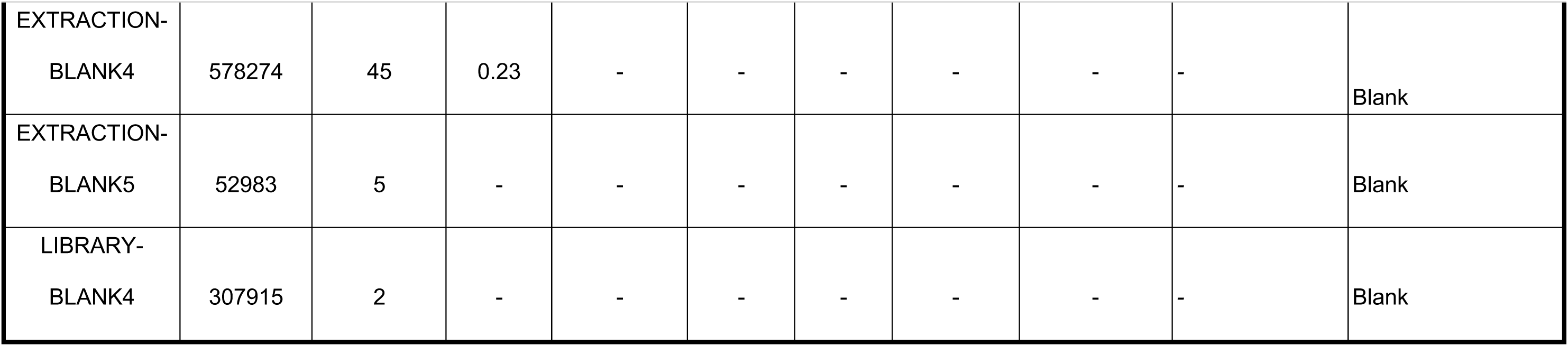
Sequencing results.

| ID | Sequenced reads | Human aligned and filtered | Human depth (x) | Animal aligned and filtered | Animal depth (x) | Damage (3') | Human: animal proportion | Proportion of target animal reads in run | Species assessment | Overall assessment |
| --- | --- | --- | --- | --- | --- | --- | --- | --- | --- | --- |
| Dz15136 | 7596370 | 30117 | 144.77 | 556 | 2.67 | 0.46 | 54.17 | 0.00007 | *Bos taurus* | Possible species ID |
| Dz2724 | 3985647 | 955 | 3.93 | 21 | 0.07 | 0.02 | 45.48 | 0.00001 | *Bos taurus* | Fail |
| Dz19364 | 3486590 | 904 | 4.00 | 60 | 0.20 | 0.02 | 15.07 | 0.00002 | *Cervus elaphus* | Possible species ID |
| Dz19352 | 4894516 | 1190 | 5.00 | 164 | 0.1 | 0.08 | 7.26 | 0.00003 | *Alces alces* | Possible species ID |
| Dz15129 | 66595 | 1 | - | - | - | 0.01 | - | - | - | Fail |
| Dz19285 | 3440798 | 10240 | 46.73 | 164 | 0.87 | 0.01 | 62.44 | 0.00005 | *Sus scrofa* | Fail: Implausible species ID; must be animal glue contaminant |
| ML4529 | 3594736 | 4592 | 18.14 | 181 | 0.8 | 0.02 | 25.37 | 0.00005 | *Sus scrofa* | Fail: Implausible species ID; must be animal glue contaminant |
| ML4530 | 5974246 | 741 | 2.76 | 58 | 0.17 | 0.02 | 12.78 | 0.00001 | *Bos taurus* | Fail: Implausible species ID; must be animal glue contaminant |
| ML4533 | 5223363 | 4454 | 16.62 | 139 | 0.47 | 0.03 | 32.04 | 0.00003 | *Bos taurus* | Fail: Implausible species ID; must be animal glue contaminant |
| ML4534 | 4144995 | 28306 | 134.02 | 892 | 2.97 | 0.03 | 31.73 | 0.00022 | *Bos taurus* | Fail: Implausible species ID; must be animal glue<br>contaminant |
| ML4532 | 3737677 | 22929 | 94.54 | 181 | 0.69 | 0.01 | 126.68 | 0.00005 | *Capra hircus* | Fail: Implausible<br>species ID; must be<br>animal glue<br>contaminant |
| Poi1 | 5406771 | 135 | 1.70 | - | - | 0.01 | - | - | - | Fail |
| Poi2 | 8950423 | 164 | 0.63 | - | - | 0.00 | - | - | - | Fail |
| LQ10 | 4177141 | 135 | 0.57 | - | - | 0.00 | - | - | - | Fail |
| Fe4 | 6870813 | 1843 | 8.55 | - | - | 0.01 | - | - | - | Fail |
| IST4 | 1269927 | - | - | - | - | 0.10 | - | - | - | Fail |
| St755 | 13668029 | 532 | 2.43 | - | - | 0.00 | - | - | - | Fail |
| St766 | 5702910 | 112 | 0.49 | - | - | 0.00 | - | - | - | Fail |
| St766b | 4071809 | 138 | 0.69 | 30 | 0.14 | 0.09 | 4.60 | 0.00001 | *Bos taurus* | Fail: Implausible<br>species ID; must be<br>animal glue<br>contaminant |
| St1017 | 4699441 | 77 | 0.35 | - | - | 0.20 | - | - | - | Fail |
| St673_powder | 8476281 | 126 | 1.00 | 1501 | 5.4 | 0.40 | 0.08 | 0.00018 | *C. elaphus* | Positive species ID |
| St784 | 8019576 | 877 | 3.50 | - | - | - | - | - | - | Fail |
| St694 | 13780889 | 12443 | 55.3 | 779 | 2.5 | 0.35 | 15.97 | 0.00006 | *B. bonasus* | Possible species ID |
| St801 | 10825776 | 30482 | 172.00 | 2976 | 11 | 0.44 | 10.24 | 0.00027 | *C. hircus* | Possible species ID |
| Dz13771 | 6965180 | 2441 | 10.70 | 1398 | 10.78 | 0.42 | 1.75 | 0.00020 | *C. hircus* | Possible species ID |
| Dz12076 | 7460673 | 13767 | 65.4 | - | - | - | - | - | - | Fail |
| Dz19307 | 7050983 | 9560 | 42.00 | - | - | - | - | - | - | Fail |
| Samele Klde | 8555243 | 313 | 0.97 | 83 | 0.26 | - | 3.77 | 0.00001 | *C. elaphus* | Possible species ID |
| CHU1 | 4444816 | 4204 | 0.69 | 3 | - | 0.12 | 1401.33 | 0.00000 | - | Fail |
| CHU2 | 4340206 | 10742 | 0.75 | 2200 | 8.222 | 0.39 | 4.88 | 0.00051 | *Cervus elaphus* | Possible species ID |
| CHU3 | 5786327 | 30464 | 1.11 | 1876 | 6.65 | 0.25 | 16.24 | 0.00032 | *Cervus elaphus* | Possible species ID |
| CHU4 | 1367638 | 2497 | 0.49 | 29371 | 176.91 | 0.34 | 0.09 | 0.02148 | *Cervus elaphus* | Possible species ID |
| KS3;<br>RGM.1333607 | 4854073 | 255 | 14.75 | 0 | - | - | - | - | - | Fail |
| KS6;<br>RGM.1333610 | 4274500 | 2278 | 6.37 | 1 | - | 0.01 | 2278.00 | 0.00000 | - | Fail |
| NR1 | 5041283 | 1262 | 5.94 | 0 | - | 0.01 | - | - | - | Fail |
| NR2 | 4737747 | 3439 | 2.19 | 0 | - | 0.01 | - | - | - | Fail |
| StEx1 | 5737159 | 14018 | - | 33 | - | 0.18 | 424.79 | 0.00001 | - | Fail |
| StEx2 | 4605353 | 557 | - | 17 | - | 0.23 | 32.76 | 0.00000 | - | Fail |
| StEx3 | 5015779 | 1504 | 12.33 | 1138 | 3.51 | 0.33 | 1.32 | 0.00023 | *Bison bonasus* | Possible species ID |
| TB1 | 157358 | 184 | 0.00 | 147 | 0.56 | 0.00 | 1.25 | 0.00093 | *Cervus elaphus* | Possible species ID |
| TB2 | 33254 | 72 | 0.16 | 13904 | 46.60 | 0.00 | 0.01 | 0.41812 | *Cervus elaphus* | Possible species ID |
| TB3 | 3886632 | 196 | 1.86 | 9708 | 33.77 | 0.00 | 0.02 | 0.00250 | *Cervus elaphus* | Possible species ID |
| EXTRACTION-BLANK1 | 196250 | 40 | - | - | - | 0.40 | - | - | - | Blank |
| LIBRARY-BLANK1 | 66595 | 1 | - | - | - | - | - | - | - | Blank |
| EXTRACTION-BLANK2 | 285807 | 39 | - | - | - | - | - | - | - | Blank |
| LIBRARY-BLANK2 | 278834 | 2 | - | - | - | - | - | - | - | Blank |
| EXTRACTION-BLANK3 | 941044 | 18 | - | - | - | - | - | - | - | Blank |
| LIBRARY-BLANK3 | 620282 | 8 | - | - | - | - | - | - | - | Blank |
| EXTRACTION-BLANK4 | 578274 | 45 | 0.23 | - | - | - | - | - | - | Blank |
| EXTRACTION-BLANK5 | 52983 | 5 | - | - | - | - | - | - | - | Blank |
| LIBRARY-BLANK4 | 307915 | 2 | - | - | - | - | - | - | - | Blank |

### Phylogenetic analyses

We constructed consensus sequences of the *Cervus elaphus* mitochondrial genome using the endoCaller function from Schmutzi (Renaud et al., 2015) and using the sequence NC_007704.2 as the reference genome. The consensus sequence (named CHU4-Chufín) was aligned with other modern and ancient *Cervidae* mtDNA (Mackiewicz et al., 2022; Rey-Iglesia et al., 2017), and the phylogenetic relationships were examined with a ML-tree (figure tree). We observe that CHU4 clusters with other samples from Liñares Cave (38,000 BP, Northern Iberia) and Holocene samples from Denmark and modern Cervus (represented by the Polish WEST1 and WEST2). All these separate from another clade only constituted by some other Liñares Cave individuals, as previously described in (Rey-Iglesia et al., 2017). These results suggest that the Chufín cervid belongs to the western lineage that split from the eastern lineage one Million years ago (Mackiewicz et al., 2022), belonging to the dominant clade encompassing European red-deer diversity (fig. 1).

**Figure 1:**
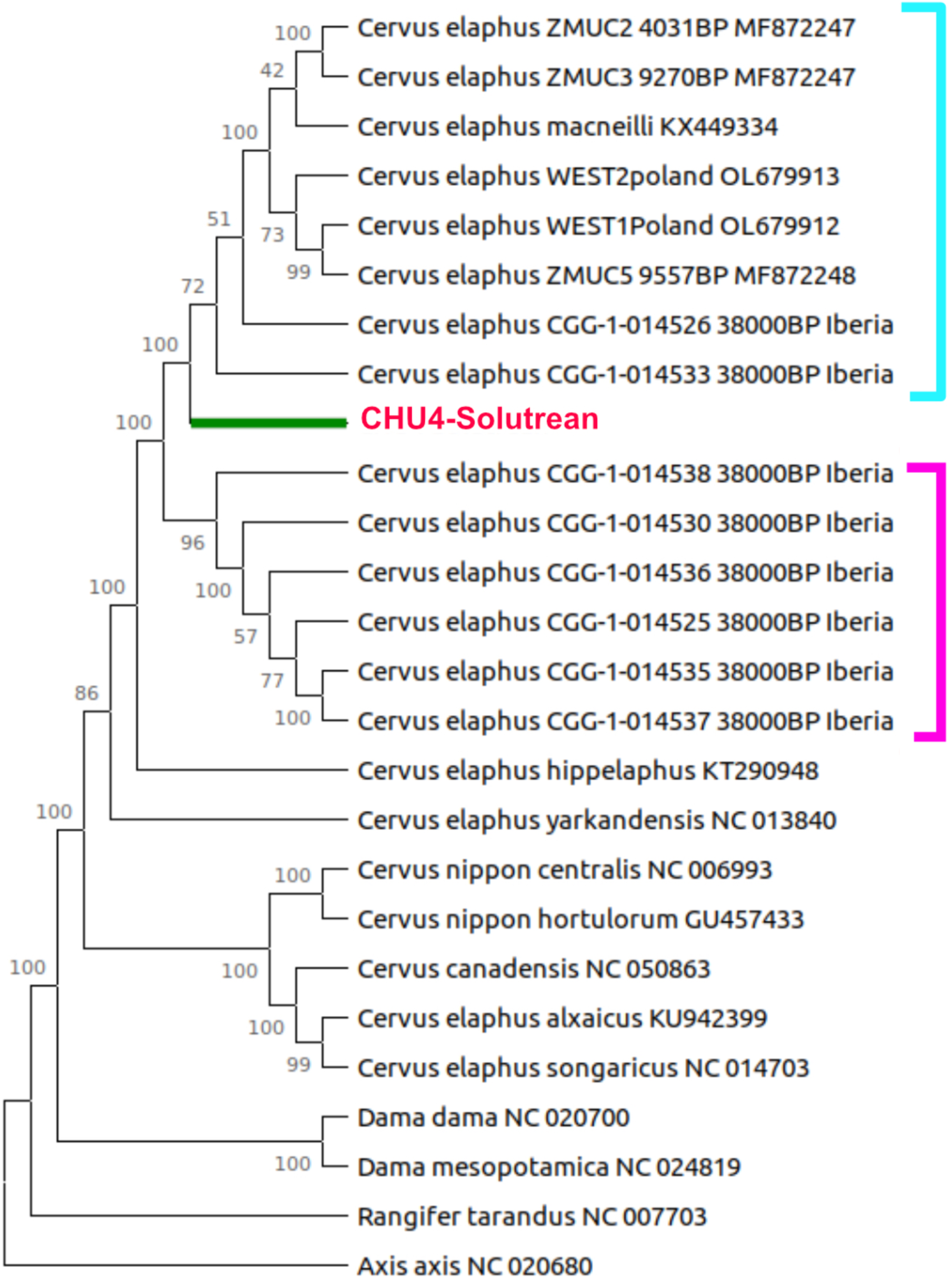
Maximum likelihood phylogenetic tree contextualising CHU4 within the Western red-deer individuals (red). The purple bracket groups the samples that Rey-Iglesia et al. (2017) define as red deer carrying foreign haplotypes associated with eastern red deer diversity. Node numbers depict bootstrap values after 100 replications.

Out of the 23 antler items tested, 8 (35%) gave enough results for a positive taxon identification. This is on par with the bone samples where 8 out of 17 (47%) were identified positively, thereby confirming antler as suitable for aDNA preservation.

For the five samples investigated for which extracts were obtained using both the traditional powdering method and the minimally invasive method (table 3), species identification was possible in two cases. Although the powdering method yielded a higher number of reads and a consequently higher coverage, failure to identify species with one method correlates with a failure of the other. It may therefore be advantageous to use the powdering method in borderline samples, but the minimally invasive method seems to perform well enough when aDNA is preserved.

**Table 3:** Comparison of minimally-invasive method with the traditional drilling.

| ID | Extraction method | Sequenced reads | Human aligned and filtered | Human depth (x) | Animal aligned and filtered | Animal depth (x) | Damage (3') | Human:animal reads proportion | Species assessment | Overall assessment |
| --- | --- | --- | --- | --- | --- | --- | --- | --- | --- | --- |
| St784 | Powder | 4628525 | 134 | 0.48 | - | - | - | - | - | Fail |
|  | Minimally invasive | 8019576 | 877 | 3.50 | - | - | - | - | - | Fail |
| St694 | Powder | 8210125 | 300 | 1.15 | 1534 | 5.60 | 0.35 | 0.1956 | *B. bonasus* | Positive species ID |
|  | Minimally invasive | 13780889 | 12443 | 55.3 | 779 | 2.50 | 0.35 | 15.9730 | *B. bonasus* | Positive species ID |
| Dz12076 | Powder | 9203358 | 1480 | 6.20 | - | - | - | - | - | Fail |
|  | Minimally invasive | 7460673 | 13767 | 65.4 | - | - | - | - | - | Fail |
| Dz19307 | Powder | 10220258 | 582 | 2.50 | - | - | - | - | - | Fail |
|  | Minimally invasive | 7050983 | 9560 | 42.00 | - | - | - | - | - | Fail |
| Samele Klde | Powder | 10147591 | 9595 | 45.8 | 1332 | 5.10 | 0.17 | 7.2035 | *C. elaphus* | Positive species ID |
|  | Minimally invasive | 8555243 | 313 | 0.97 | 83 | 0.26 | - | 3.7711 | *C. elaphus* | Positive species ID |

As part of the method improvement, a 20-minute pre-digestion step was performed on some of the extracted items. This step demonstrated clear benefits by reducing the proportion of human contaminant reads compared to endogenous animal reads, as this is improved in all but one of the cases that did not fail altogether (table 4).

**Table 4:** Assessment of the effect of the predigestion process.

| ID | EXTRACT |  |  |  |  |  | PREDIGEST |  |  |  |  |  | Comparison |
| --- | --- | --- | --- | --- | --- | --- | --- | --- | --- | --- | --- | --- | --- |
|  | Sequenced reads | Human aligned and filtered | Animal aligned and filtered | Damage | Human:animal reads | Species assessment | Sequenced reads | Human aligned and filtered | Animal aligned and filtered | Damage | Human:animal reads | Species assessment |  |
| CHU1 | 191 | 177 | 3 | 0.12 | 59.0000 | - | 13680721 | 88 | - | 0.16 | - | - | -- |
| CHU2 | 1550 | 181 | 2200 | 0.39 | 0.0823 | *Cervus elaphus* | 11504150 | 296 | 1138 | 0.35 | 0.2601 | *Cervus elaphus* | Improvement after predigestion |
| CHU3 | 1213 | 289 | 1876 | 0.25 | 0.1541 | *Cervus elaphus* | 30577831 | 119 | 170 | 0.21 | 0.7000 | *Cervus elaphus* | Improvement after predigestion |
| CHU4 | 30242 | 121 | 29371 | 0.34 | 0.0041 | *Cervus elaphus* | 13249226 | 341 | 11606 | 0.20 | 0.0294 | *Cervus elaphus* | Improvement after predigestion |
| KS3 | 28 | 1260 | 0 | 0.00 | - | - | 12007941 | 1288 | - | 0.00 | - | - | -- |
| KS6 | 371 | 3419 | 1 | 0.01 | 3419.0000 | - | 12221938 | 3345 | - | 0.01 | - | - | -- |
| NR1 | 192 | 1489 | 0 | 0.01 | - | - | 13128526 | 1423 | - | 0.03 | - | - | -- |
| NR2 | 46 | 543 | 0 | 0.01 | - | - | 10383084 | 191 | 116 | 0.17 | 1.6466 | *Cervus elaphus* | Extraction failed but information in predigest |
| StEx1 | 7 | 18 | 33 | 0.18 | 0.5455 | - | 12703363 | 244 | - | 0.12 | - | - | -- |
| StEx2 | 37392 | 72 | 17 | 0.23 | 4.2353 | - | 9065039 | 295 | - | 0.18 | - | - | -- |
| StEx3 | 43375 | 3006 | 1138 | 0.33 | 2.6415 | *Bison bonasus* | 11032868 | 489 | 198 | 0.14 | 2.4697 | *Bison bonasus* | Improvement after predigestion |
| TB1 | 75 | 17 | 147 | 0.00 | 0.1156 | *Cervus elaphus* | 34172 | 290 | 217 | 0.00 | 1.3364 | *Cervus elaphus* | Less animal reads in final extract, but lower proportion of human |
| TB2 | 17096 | 38 | 13904 | 0.00 | 0.0027 | *Cervus elaphus* | 121217 | 5 | - | 0.35 | - | - | Improvement after predigestion |
| TB3 | 5786 | 543 | 9708 | 0.00 | 0.0559 | *Cervus elaphus* | 20850 | 18 | - | 0.00 | - | - | Improvement after predigestion |

From a visual macro analysis, we could document the identical marks before and after the extraction, thus allowing us to reach the same conclusion regarding the technological considerations (reconstitution of the operational sequence of manufacture and use, by functional macro-fractures, of the items).

Using µCT imaging, we assessed the impact of the minimally invasive extraction on the integrity of the items. According to further 3D analyses of one of the items, scanned at a resolution of 23 µm, the average erosion of the surface is between 0 and 0.200 mm. Only in very limited areas, the maximum surface erosion caused by our approach reaches 0.300 mm. Red spots visible in Figure 2 represent areas with material loss of more than 1 mm, thus areas where the extraction process caused sediment removal. In addition, 14 of the items were closely examined with a 3D microscope before and after extraction. Technologically critical marks remained visible (fig. 3).

**Figure 2:**
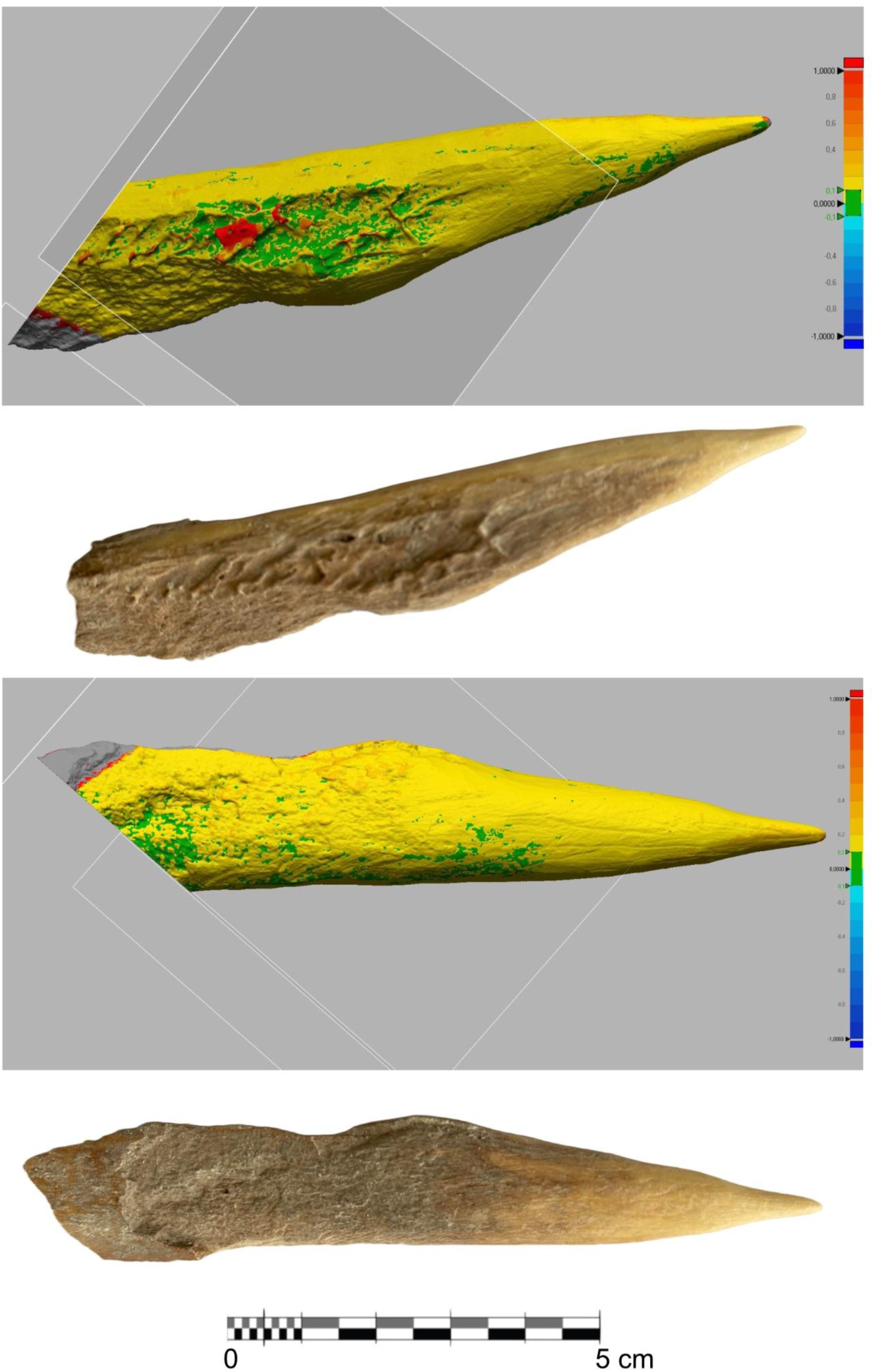
Micro-CT scan of StEx1 after sampling. Surfaces coloured in green are unmodified. The yellow colour shows a slight surface modification between 0.100-0.300 mm depth, resulting from adhered sediment removal.

**Figure 3:**
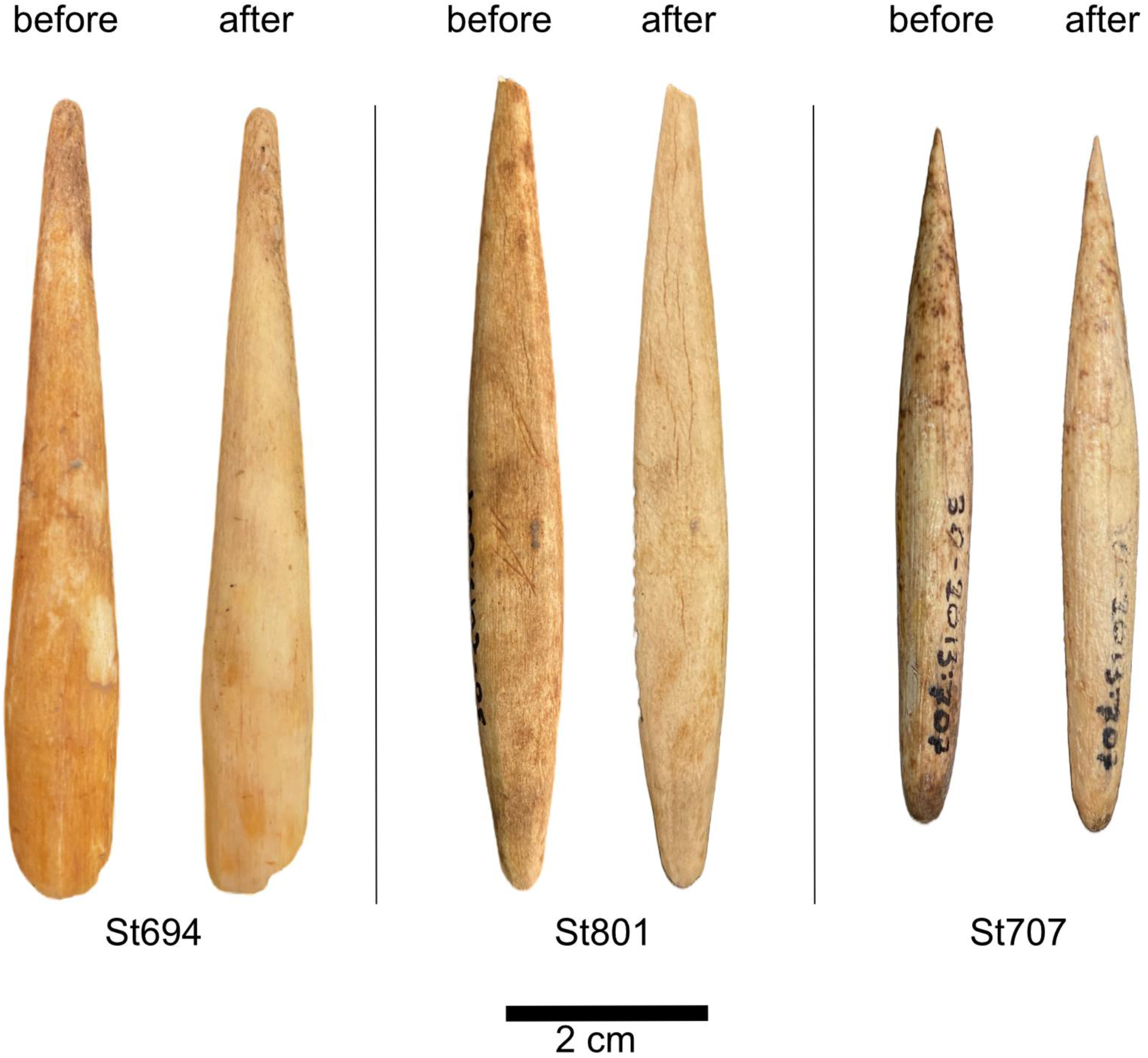
Hunting weapons (projectile points) from Satsurblia (Georgia) before and after sampling.

**Figure 4:**
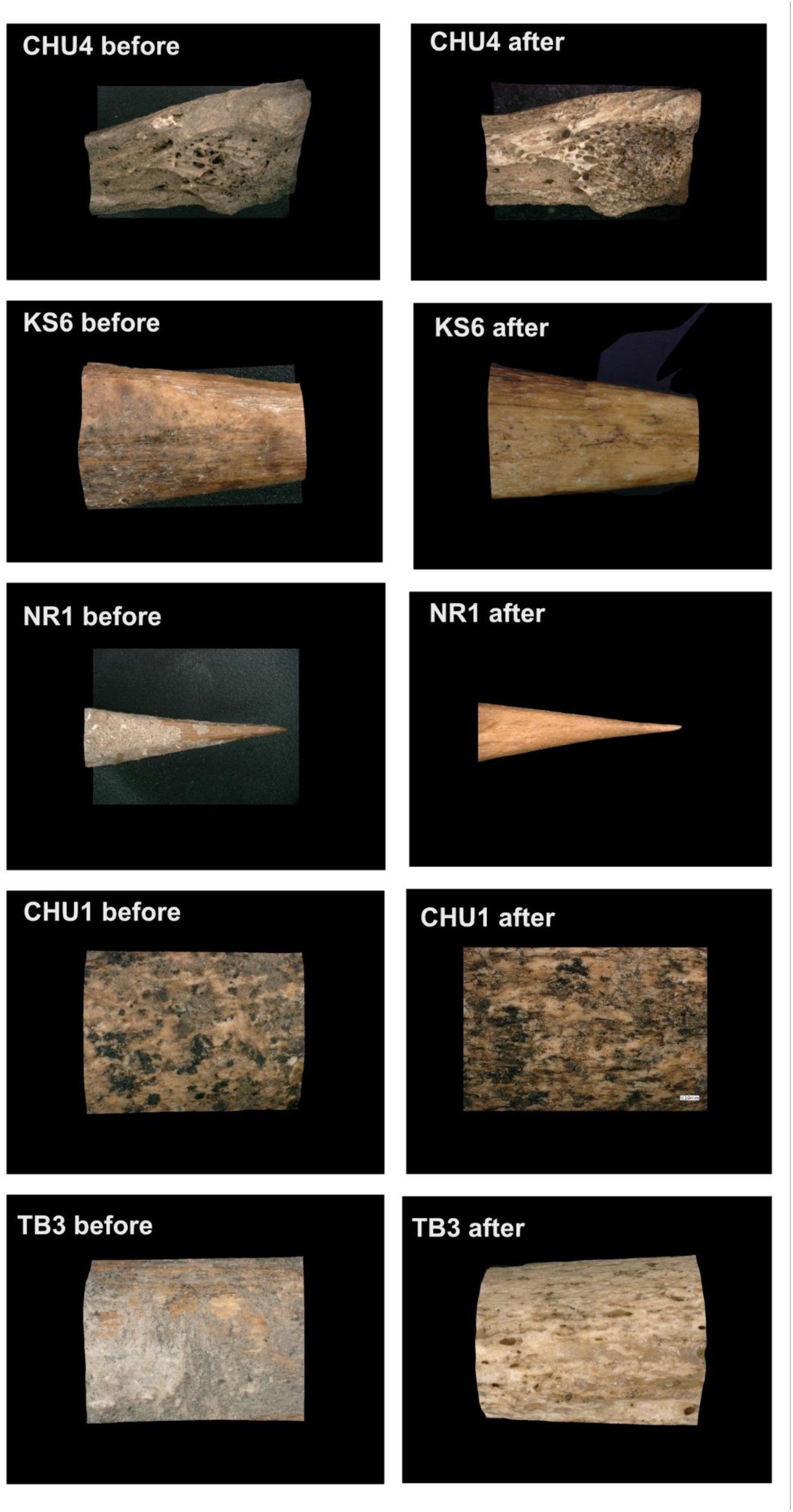
3D microscope surface images showing the effect on the sampled items’ surfaces, mainly consisting of removing the adhered sediments.

## Discussion

The obtained results indicate that it is possible to extract ancient endogenous DNA from some prehistoric bone and antler objects, applying a minimally destructive protocol. This is particularly relevant for studying the Palaeolithic osseous industry, as representative items are relatively rare, each being unique in its own way, and frequently, but not exclusively, made from antler raw materials.

Aside from macroscopic or microscopic evaluations (Lefebvre et al., 2016; Pacher, 2010), species identification of the raw material used to produce osseous artefacts can be achieved through biomolecular methods. Paleoproteomics is an approach currently favoured for this type of work. ZooMs characterises collagen proteins using peptide mass-fingerprinting (Brandt et al., 2018; Buckley et al., 2009; Collins et al., 2010). These analyses are relatively cheap and require little input material making them quasi-non-destructive (McGrath et al., 2019). However, this approach has a number of limitations in the context of bone tool studies. At present, the correct assignment of a given tool to a specific cervid species remains difficult, if not impossible (Brandt et al., 2018; Buckley et al., 2009; von Holstein et al., 2014). This is a major limitation as a high proportion of bone tools, especially Eurasian Upper Palaeolithic hunting weapons (but also some domestic tools, personal ornaments and mobile art) were produced from antlers (Averbouh, 2000; Goutas, 2004; Knecht, 1991; Tejero, 2014). Furthermore, although ZooMs may be able to identify the species (or family) of animals exploited for the production of an item, this method cannot determine the sex of the animal and produce phylogenetically information, unlike genomic data.

Ancient DNA, on the other hand, can provide unique information. Under certain conditions (good preservation of aDNA and absence of modern contamination), the DNA of the craftsman and/or users of the antler objects could be extracted from the object surface manipulated by prehistoric people. When reaching the target, hunting weapons are entirely impregnated with blood and other organic tissues (hair, skin, muscles, tendons, etc.) (Pétillon, 2022). Thus, it is theoretically possible to sequence and identify the DNA of both the animal exploited to produce the projectile point, and that of the prey it has been used to hunt.

To extract ancient DNA, traditional methods involve the drilling of powder, which is then dissolved to release the DNA trapped in the mineral and organic fractions of the bone (e.g., (Dabney et al., 2013) protocol). However, by modifying the approach to avoid the necessity of producing a powder, it becomes possible to release endogenous DNA without significantly affecting the item’s integrity. This approach was explored by Harney et al. (2021) for human teeth. One fundamental element of their suggested protocol involved the protective wrapping of the teeth to ensure that only a small part of it is exposed to the DNA extraction. This aspect had to be modified for the study of bone tools for two important reasons. Firstly, the larger nature of many of these items required the use of larger amounts of extraction buffer, resulting in very dilute lysates if only a small area of the piece were to be exposed. In addition, whereas teeth have a very standard structure, with the cellular cementum as a main target of the extraction (Harney et al., 2021), bone tools are much less standardised with no *a priori,* obviously DNA-richer zones. Furthermore, exposing the entire piece to extraction buffer enabled the extraction from a larger surface area, thereby releasing more DNA. This is of double importance in the context of osseous industry studies. Indeed, much of this work is focused on observing the technical marks on the pieces’ surfaces, which remain visible using this approach. In addition, due to their scarcity and appearance, osseous industry elements are frequently displayed in museum contexts. By performing a treatment that affects the entire piece in a uniform manner, as is the case here, the items’ aesthetic character is preserved.

The numerous positive results obtained confirm that the decontamination protocol performed chemically removing potential surface contamination using bleach and short-wave UV light for these minimally-invasive extractions was adequate. Although some modern human contamination does remain, it does not overwhelm animal reads to the point of no longer being able to identify them. It nevertheless remains clear that the conservation and handling of the objects play a fundamental role in the amount of contamination detected. Most of the objects in this study were subjected to some level of conservation and handling. In contrast, those from the two Spanish sites (Cueva Chufín and Tito Bustillo) were collected during excavation, with the intention of aDNA studies, and were not handled with bare hands. This is reflected in the obtained results, as, with the exception of one failed sample (CHU1), all had more animal than human reads.

For the objects from Mladeč, we obtained mammalian mitochondrial data. However, considering the previous visual observation, the taxonomic attribution was entirely implausible. Four out of the five items were made from antler, and one (ML4530) was manufactured from ivory. However, the mitochondrial DNA found was attributed to *Sus scrofa*, *Bos taurus* and *Capra hircus*, three common domesticates. As these items were excavated more than 100 years ago, collected by J. Knies (Oliva, 2006), and subsequently conserved in the Moravian Museum, it is likely that, through time, animal-based glues and consolidants may have been applied to these items, and those are the ones identified by our analyses (Schellmann, 2007). Unfortunately, due to a common lack of record keeping of conservation measures applied to pieces in older collections, this issue is likely to be frequent when working with material excavated more than a few decades ago. It therefore highlights the necessity to combine multiple analyses of individual items to maximise knowledge gained about them as well as cross-validate obtained results.

Although positive species identifications have been achieved using this minimally-invasive method, some limitations remain. The amount of obtained data is very small. Only for one of these specimens, has it been possible to obtain enough data to perform more detailed analyses to identify its taxonomy. In addition to the methodological challenges associated with genetic studies of osseous industry material, it is also important to note that the majority of the objects studied in the above-presented work were made from antler. Although similar to bone in many respects, antler is a distinct material, which has not been a substrate of choice for the retrieval of aDNA. Antlers are an exoskeletal appendage characteristic of the *Cervidae* (deer) family with a yearly cycle of growth, fall and regrowth. They comprise bone, cartilage, fibrous tissue, skin, nerves, and blood vessels and are generally found only in males, except for reindeer (*Rangifer tarandus*) (Crigel et al., 2001; Goss, 1983; Kierdorf & Kierdorf, 2004). Only one study has retrieved ancient DNA from palaeontological antlers from the Allerød interstitial period (c. 12,000 years) (Kuehn et al., 2005), and only two from recent, historical contexts (Rosvold et al., 2019; von Holstein et al., 2014). We therefore confirm that aDNA can be successfully retrieved from Pleistocene antler material. Furthermore, just like bone, the taphonomic history of the objects is of fundamental importance to the long-term DNA preservation in the items, more so that their absolute age.

To assess potentially adverse effects of the here-presented minimally-invasive method on the sampled items, a combination of visual monitoring during the sampling and a 3D microscopic surface analysis and micro-CT of the surface and the inner piece before and after sampling were performed.

The destructive sampling process associated with the aDNA analyses is problematic for unique pieces, as can be the case for the Palaeolithic objects in osseous materials. Recent studies stress the importance of evaluating the potential effects on bone tools preservations of the non-destructive or minimally invasive techniques (Martisius, Welker, et al., 2020; Mateo-Lomba et al., 2022; Sinet-Mathiot et al., 2021). Our study demonstrates that this minimally-invasive method can successfully be applied.

## Conclusion

Our results demonstrate that antler objects can be a source of aDNA. While paleogenomics focuses on bone and teeth as the primary tissues to obtain aDNA, we add another possible source. Given the importance of antlers as a raw material for the hunter-gatherer groups at the end of the Pleistocene, but also for later societies until medieval times, it is critical to obtain as much data as we can by combining archaeological and biomolecular methods. Moreover, by applying a minimally-invasive method on bone and antler objects based on (Harney et al., 2021), we contribute to preserving the integrity of such archaeologically unique items. To this day, the oldest antlers to yield aDNA come from palaeontological contexts of around 12ka (Kuehn et al., 2005). Our study shows that antlers older than 30ka could also be a reliable source of aDNA. This provides another tool to deepen our knowledge of Upper Palaeolithic societies from the earlier *H. sapiens* permanently established in Eurasia to recent Prehistoric times.

## Acknowledgements

The authors would like to thank Maayan Shemer, Nimrod Marom, and Omry Barzilai for providing access and assisting in researching the material from Nahal Rahaf 2 Rockshelter, as well as Jürgen Kriwet and Sebastian Stumpf for access to the 3D microscope. Thanks are due to Natasja den Ouden for giving us access to the material of Kars Akil stored at the Naturalis Biodiversity Center (NL). We are grateful to the following institutions for access to the samples: the Musée d’Archéologie Nationale (Saint Germain-en-Laye. France. Items from Isturitz, La Quina-Aval, La Ferrassie), The Anthropos Institut of the Moravian Museum (Brno. Czech Republic. Items from Mladeč); The Naturalis Biodiversity Center (Leiden. The Netherlands. Items from Ksâr ‘Akil); The University of Salamanca (Spain. Items from Tito Bustillo); The University of Cantabria (Santander. Spain. Items from Cueva Chufín); The University of Barcelona (Spain. Items from Cova del Parco); and the Georgian National Museum (Tbilisi. Georgia. Items from Satsurblia and Semele Klde).

## Funding

Research of J.-M. T. is supported by a project of the Meitner Program of the Austrian Science Fund (FWF) (Project: *Osseous Hunting Weapons of Early Modern Humans in Eurasia.* Number M3112). The whole project was supported by the University of Vienna Research Platform: Mineralogical Preservation of the Human Biome from the Depth of Time (MINERVA). J.-M. T., P. G., and O. C. benefited from a Seed Grant from the HEAS (Human Evolution and Archaeological Sciences) of the University of Vienna (Project: *Assessing the differential DNA preservation in Palaeolithic sediments and osseous tools from museum collections*). D. M. B. supported by a Seal of Excellence Fellowship of the Austrian Academy of Sciences (‘TechnoBeads’ project no. 101061287). P. R. N. benefited from funding by the University of Vienna and the Land Niederösterreich, Abteilung Wissenschaft & Forschung (project K3-F-530/005-2021).

### Author contributions

Conceptualization: JMT, OC, PG, RP

Methodology: JMT, OC, PG, RP

Investigation: JMT, OC, PG

Visualization: JMT, OC, PG

Funding acquisition: JMT, OC, PG, RP, MDB, PRN

Project administration: JMT

Supervision: JMT, RP

Writing – original draft: JMT, OC, PG

Writing – review & editing: JMT, OC, PG, RP, ABC, PN, MDB, PRN, AA, PGS, GW

### Data and materials availability

All the sequenced genetic data is available at ENA through the accession number: PRJEB61082

