## Supplementary table 1 for "A minimally-invasive method for ancient DNA sampling of Prehistoric bone and antler tools and hunting weapons"

Supplementary table 1: Species and sequences included in the mammalian capture kit

| NCBI code | Species |
|---|---|
| NC_020677.1 | Alces alces |
| NC_049122.1 | Apodemus sylvaticus |
| MN122828.1 | Arvicola amphibius |
| NC_014044.1 | Bison bonasus |
| NC_006853.1 | Bos taurus |
| NC_005044.2 | Capra hircus |
| NC_020684.1 | Capreolus capreolus |
| NC_028625.1 | Castor fiber |
| NC_007704.2 | Cervus elaphus |
| NC_037888.1 | Cricetus cricetus |
| NC_056768.1 | Crocidura russula |
| NC_024819.1 | Dama mesopotamica |
| NC_034307.1 | Dicrostonyx hudsonius |
| NC_005129.2 | Elephas maximus |
| MN935777.1 | Eliomys quercinus |
| NC_054160.1 | Ellobius talpinus |
| NC_001640.1 | Equus caballus |
| NC_002080.2 | Erinaceus europaeus |
| NC_001700.1 | Felis catus |
| NC_008156.1 | Galemys pyrenaicus |
| NC_020710.1 | Gazella subgutturosa |
| KM820836.1 | Rattus norvegicus |
| NC_012920.1 | Homo sapiens |
| NC_020669.1 | Hyaena hyaena |
| MT240946.1 | Hystrix brachyura |
| NC_004028.1 | Lepus europaeus |
| NC_027083.1 | Lynx lynx |
| MN935776.1 | Marmota marmota |
| NC_021749.1 | Martes martes |
| NC_011125.1 | Meles meles |
| NC_027932.1 | Micromys minutus |
| NC_038176.1 | Microtus arvalis |
| NC_005089.1 | Mus musculus |
| NC_025516.1 | Mustela erminea |
| NC_020638.1 | Mustela putorius |
| NC_025559.1 | Neomys fodiens |
| NC_001913.1 | Oryctolagus cuniculus |
| NC_001941.1 | Ovis aries |
| NC_010641.1 | Panthera pardus |
| NC_007703.1 | Rangifer tarandus |

| | |
|---|---|
| NC_001665.2 | Rattus norvegicus |
| NC_001779.1 | Rhinoceros unicornis |
| NC_020633.1 | Rupicapra rupicapra |
| NC_020746.1 | Saiga tatarica |
| NC_027963.1 | Sorex araneus |
| NC_027692.1 | Stylodipus telum |
| NC_000845.1 | Sus scrofa |
| NC_002391.1 | Talpa europaea |
| MN326850.1 | Terricola subterraneus |
| NC_003427.1 | Ursus arctos |
| NC_008434.1 V | Vulpes vulpes |
